# Pupil Constriction Causes Activity in the Human Retina and Visual System

**DOI:** 10.64898/2026.04.14.718411

**Authors:** Sebastiaan Mathôt, Olaf Dimigen, Hakan Karsilar, Veera Ruuskanen, Daria Weiden, Ana Vilotijević

## Abstract

Pupil responses shape the earliest stages of visual perception by regulating the amount of light that enters the eye. How this affects visual processing is still poorly understood. Here we present evidence that pupil constriction causes activity in the human retina and visual system, independently of visual stimulation. Healthy human participants (N=119) viewed brief visual stimuli (light increments or decrements) while pupil size, retinal activity (electroretinogram), and brain activity (electroencephalogram) were recorded. As expected, visual stimulation triggered an initial burst of retinal activity followed by pupil constriction (the pupil light response). Importantly, we used trial-to-trial variability in constriction latency to reveal a previously unknown retinal response that is locked to pupil constriction, rather than to visual stimulation. Presumably, and in line with similar findings in mice, this constriction-locked retinal activity is a response to the sudden decrease in retinal light exposure that accompanies pupil constriction (although contribution of iris muscle activity is not conclusively ruled out). A similar constriction-locked response emerged later over visual cortex. Given these findings, an important open question is how the visual system maintains brightness constancy despite pupil-induced retinal and cortical activity.

## Pupil Constriction Causes Activity in the Human Retina and Visual System

Vision starts when light touches the surface of the eye. Large pupils allow more light into the eye, which increases retinal light exposure but reduces optical quality due to greater optical aberrations. In contrast, small pupils allow less light into the eye, which decreases retinal light exposure but improves optical quality because light is more sharply focused (1). Through these effects, pupil size affects the earliest stages of visual processing.

Pupil size is typically treated as a factor that *modulates* how the visual system responds to external stimuli (reviewed in 2). Evidence for this modulatory role is observed at multiple levels of the visual system. At the retinal level, visual stimuli elicit responses that can be measured with electroretinography (ERG), using electrodes placed on either the cornea or the eyelids. When the pupil is pharmacologically dilated, the amplitude of ERG responses to visual stimuli increases (3–5). At the cortical level, visual stimuli elicit event-related potentials (ERP) that can be measured with electrodes placed on the skull. When pupils are large, either due to an experimental manipulation or spontaneous fluctuations, ERPs are affected—though the pattern is more complex than the simple amplitude increase observed for retinal responses (6–8). Finally, pupil size affects behavioral performance, such that small pupils are associated with enhanced discrimination of fine detail (visual acuity) whereas large pupils are associated with enhanced detection of faint stimuli (visual sensitivity; 9–15). Taken together, pupil size modulates processing of external stimuli at all levels of the visual system. However, even in the absence of external stimuli (and except in complete darkness), changes in pupil size result in changes in retinal light exposure. Could these pupil-driven changes in retinal input trigger responses in the visual system *as if* there was an external stimulus? In other words, might pupils not only modulate visual processing but also *generate* activity within the visual system? Lapanja and colleagues (16) measured activity of retinal ganglion cells (RGCs) in living mice to test whether the mouse retina responds to pupil-constriction-induced stimulation. They presented brief flashes of light to one eye, which triggered pupil constriction in both eyes (because the pupil light response is consensual). Crucially, they observed RGC activity in the non-stimulated eye that coincided with pupil constriction. This activity was presumably driven by the sudden decrease in retinal light exposure caused by the constricting pupil. The results of Lapanja and colleagues (16) thus suggest that pupil constriction indeed triggers retinal responses, even in the absence of external stimuli.

A follow-up experiment suggested that a similar mechanism may be present in humans (16). When viewing a sustained full-field flash, observers reported a perceptual dimming of the stimulus around the time of pupil constriction, despite no actual luminance change. However, human retinal activity was not recorded, leaving open the question of whether the perceptual effect was indeed due to pupil-constriction-induced retinal activity, or whether it reflected a perceptual after effect unrelated to pupil size.

Here we set out to replicate and extend these important results by investigating whether a pupil-constriction-induced response exists in the human retina and visual system. To do so, we conducted a combined analysis of several large datasets collected in our lab. In all datasets, participants (N=119) were exposed to brief visual stimuli while pupil size, retinal activity (ERG), and cortical activity (ERP) were recorded. As expected, stimuli triggered an initial burst of retinal activity followed by pupil constriction (the pupil light response). We leveraged trial-to-trial variability in pupil-constriction latency to dissociate stimulus-locked from pupil-constriction-locked ERG and ERP responses. To foreshadow the results, we found a previously unknown ERG response that a) is locked to pupil constriction, b) originates from the retina, c) scales with pupil-constriction velocity, and d) is robustly observed under a wide range of conditions. In addition, we found a small constriction-locked ERP that emerged over occipital electrodes around 100 ms after the constriction-locked ERG response, plausibly reflecting a pupil-constriction-locked response in early visual cortex. These findings provide important new insights into how changes in pupil size affect the human retina and visual system, and the role of pupil size in visual perception more generally.

## Methods

### Open-practices statement

All data, experimental materials, analysis scripts, and *Supplementary Materials* are available from https://osf.io/2u6jv/. (Since this manuscript presents a combined analysis from various datasets, the data has been preprocessed and merged into a single structure.) The study was not preregistered. We therefore emphasize general patterns over significance of individual effects.

### Datasets

This is an analysis of nine separate datasets with a total of 119 participants, all healthy, adult human participants with normal, uncorrected vision. Datasets were collected for a different purpose. Details, including stimulation parameters, are described in the *Supplementary Materials*. In all datasets, participants viewed a brief visual stimulus that triggered both a retinal response and a pupil constriction.

### Materials and software

All datasets were recorded in the same laboratory with similar software and under similar conditions. The experimental scripts were implemented using OpenSesame 4.0 (17) using the PsychoPy (18) backend for display presentation and PyGaze (19) for eye tracking.

The experiments were conducted on a desktop computer with a 27” LCD monitor with a refresh rate of 120 Hz and a resolution of 1920 × 1080 pixels. At the viewing distance of 76 cm, the full-screen stimuli subtended 44.8° x 25.2° degrees of visual angle. Pupil diameter and gaze position were recorded from the right eye at 1000 Hz using an EyeLink 1000 (SR Research) eye tracker. For datasets with EEG, data was recorded at 1000 Hz from 26 electrodes placed according to the standard 10-20 system using a TMSI REFA 32 amplifier controlled by OpenViBE data acquisition software (20). Two additional electrodes were placed on the left and right mastoids for offline re-referencing. Two additional channels were placed immediately above the eyelids and on the infraorbital margin of each eye (four channels in total; Fig. 1). During recording, all electrodes (EEG and ERG) were referenced against the average of all electrodes, and then re-referenced offline against the average of the left and right mastoids. After preprocessing, all data was downsampled to 250 Hz.

**Figure 1.**
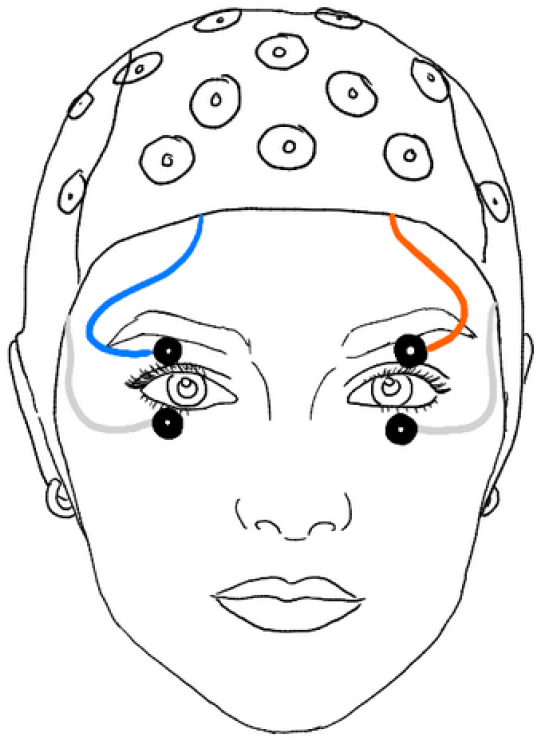
EOG electrodes were placed above and below each eye. The ERG signal corresponds to the average of those four channels.

Data analysis was performed using the Python `eeg_eyetracking_parser` (6) toolbox, which is a high-level toolbox that uses `MNE` (21) for general EEG processing, `eyelinkparser` (22) for eye-movement and pupil-size processing, `autoreject` (23) for rejecting and repairing bad EEG channels and epochs, and `mne-icalabel` for a control analysis to rigorously exclude contributions of eye and muscle artifacts (24).

### Data preprocessing

#### Pupil size

For the pupil-size analyses, we followed the recommendations from Mathôt and Vilotijević (22). 1) Missing or invalid data was interpolated using cubic-spline interpolation if possible, using linear interpolation if cubic-spline interpolation was not possible (when the segment of missing data was too close to the start or end of a trial), and removed if interpolation was impossible altogether (when data was missing from the start and/ or until the end of a trial) or if the period of missing data was longer than 500 ms and thus unlikely to reflect a blink (see `datamatrix.series.blinkreconstruct`). 2) Pupil size was converted from arbitrary units as recorded by the EyeLink to millimeters of diameter. No baseline correction was applied to pupil size.

#### Timepoint of maximum constriction velocity

To determine the timepoint of maximum constriction velocity, pupil size was first smoothed with a 12 ms Hanning window. Next, within a search window of 200 to 880 ms after stimulus onset, the point at which the velocity of this smoothed signal was most negative was taken as the timepoint of maximum constriction. The velocity of the smoothed signal at this moment was taken as the maximum constriction velocity.

#### Electroencephalography (EEG) and electroretinography (ERG)

EEG and ERG data was preprocessed fully automatically. Unless otherwise specified, we used the default parameters as described on the documentation of the referenced functions. 1) Data was re-referenced to the average of the mastoid channels. 2) Muscle artifacts, which are characterized by bursts of high-frequency activity, were marked as bad using the MNE function for annotating muscle activity (see `mne.preprocessing.annotate_muscle_zscore`) with a z-threshold of 5. 3) Data was filtered using a 0.1 - 40 Hz bandpass. 4) The RANSAC algorithm (see `autoreject.Ransac`) identified bad channels based only on data segments corresponding to trials; in brief, this algorithm assumes that a channel is bad if its data is poorly predicted by interpolation from neighboring channels (25). Baseline correction was applied to the ERG or EEG signal (but not to pupil size) using the 100 ms preceding stimulus onset as the baseline period. The ERG signal corresponds to the average of all four EOG channels, referenced against the mean of the left and right mastoid.

#### Trial exclusion

Trials were excluded from analysis if 1) a blink as identified by the EyeLink’s built-in algorithm occurred within 500 ms after stimulus onset, 2) prestimulus pupil size was unusually small (< 1.5 mm) or large (> 8 mm), 3) the standard deviation of the ERG signal exceeded 25 µV (this threshold was determined based on a visual inspection of the trial distribution of standard deviations), 4) the timepoint of maximum constriction was near the boundaries of the search window, indicating an unrealistically fast (< 180 ms) or slow (> 860 ms) pupil light response; or the maximum constriction velocity was unrealistically high (-.08 a.u.) or low (-.005 a.u.) based on a visual inspection of the trial distribution of constriction velocities.

Based on the above criteria, 48,509 of 55,345 trials (87.6%) remained for further analysis. Within reasonable limits, none of the reported results depend on the specific exclusion criteria.

## Results

### Key result: An ERG component locked to pupil constriction

The main finding is the existence of a previously unknown ERG component locked to the timepoint of maximum constriction velocity (Fig. 2a). It is characterized by a small initial negative deflection (in some datasets, see below), followed by a larger positive deflection (in all datasets) that roughly coincides with the timepoint of maximum constriction velocity. It is a small component—roughly an order of magnitude smaller than the a- and b-waves, which are triggered by external visual stimulation (Fig. 3). Presumably, the constriction-locked ERG component is a response to the decrease in retinal exposure that results from pupil constriction, as has previously been reported for the mouse retina (16).

**Figure 2.**
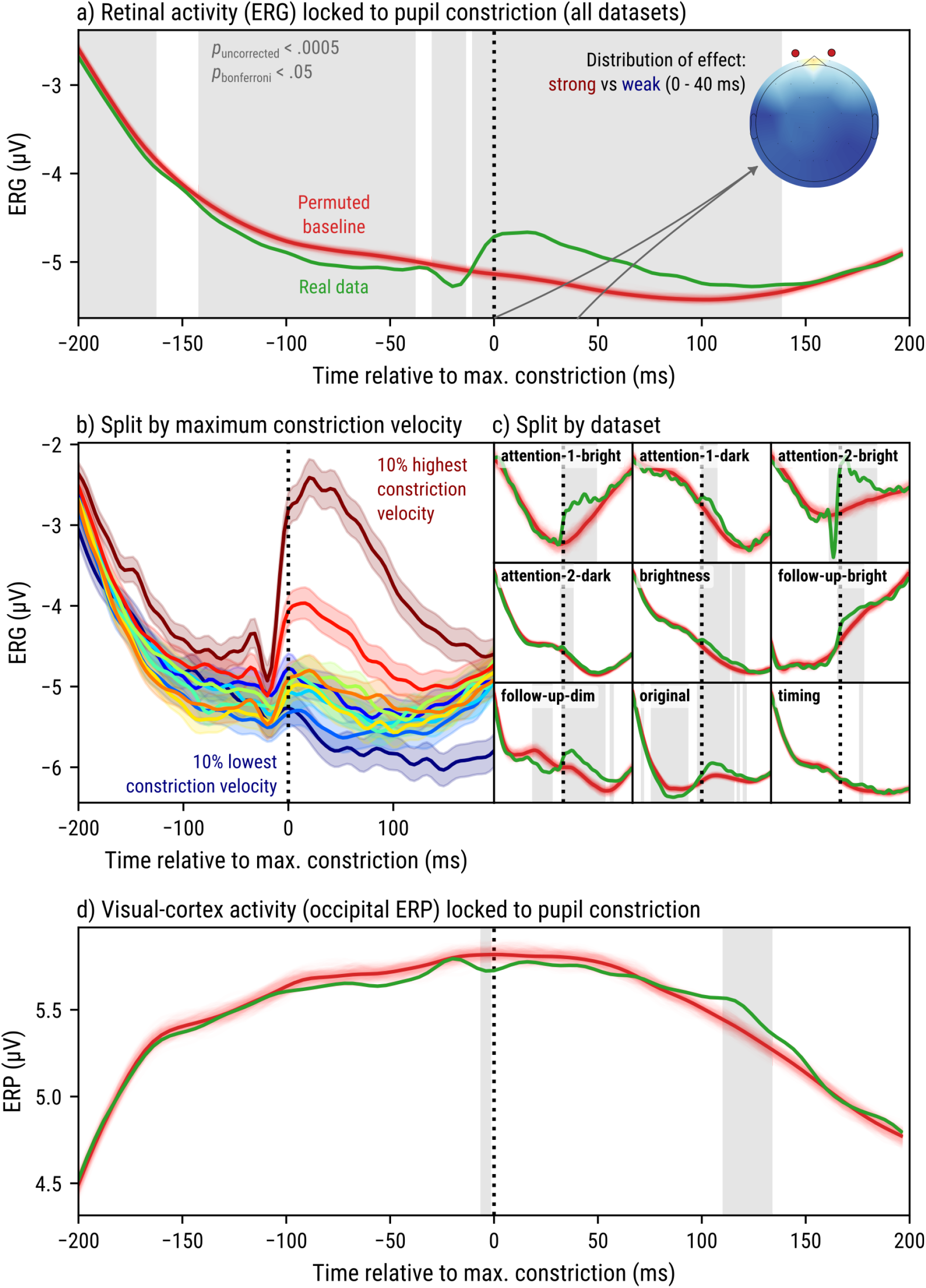
a) Retinal activity (ERG) across all datasets and locked to the timepoint of maximum constriction velocity. Green lines correspond to the actual ERG signal. Red lines correspond to permuted baselines (N = 2000). Gray shadings indicate significant differences between the actual ERG signal and permuted baselines. The inset scalp distribution illustrates the distribution of effect sizes (difference between actual ERG signal and permuted baselines) across scalp and eye electrodes. b) Constriction-locked ERG as a function of maximum constriction velocity. c) Constriction-locked ERG and permuted baselines for individual datasets. d) Visual-cortex activity (ERP averaged across occipital electrodes O1, O2, and Oz) across all datasets and locked to constriction.

**Figure 3.**
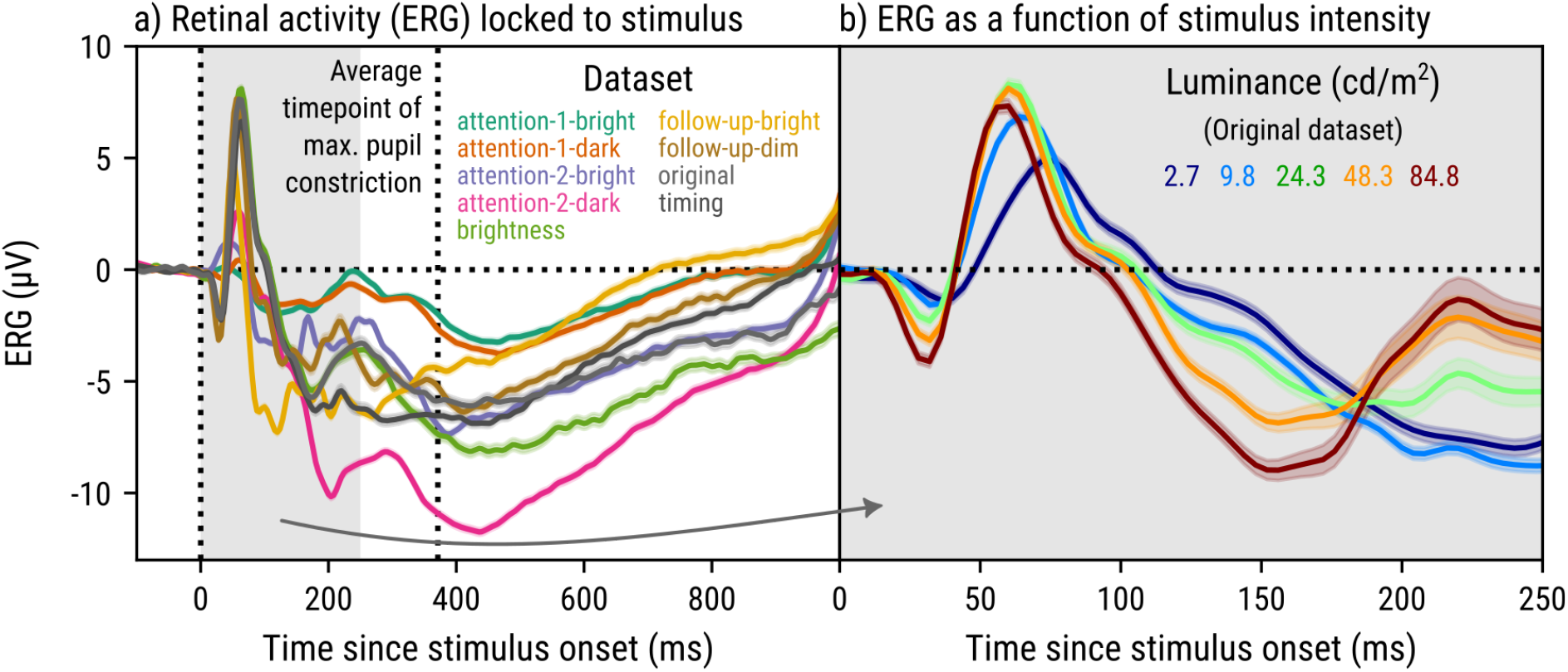
a) Retinal activity (ERG) locked to stimulus onset for all datasets. The second vertical dotted line indicates the average timepoint of maximum constriction velocity. b) ERG for the Original dataset, zoomed in on the first 250 ms and split by stimulus intensity.

#### Permutation testing confirms a constriction-locked ERG component

The constriction-locked component was tested against a permuted baseline. First, timepoints of maximum constriction velocity were shuffled within groups of trials for the same participant, dataset, and stimulation condition (i.e. each trial was assigned the timepoint from another trial). Next, the signal was locked to these shuffled timepoints, resulting in a single permuted baseline. This procedure was repeated 2000 times. Combining this procedure with a cluster-based permutation test (or other sophisticated forms of multiple-comparison correction) is extremely computationally intensive; for this reason, and because the difference between the real and shuffled signals is clearly very large, we applied highly conservative (but computationally cheap) Bonferroni correction and used an alpha level of .05 (*p*_*bonferroni*_ < .05, *p*_*uncorrected*_ < .0005).

#### Higher maximum constriction velocity is associated with a more pronounced constriction-locked ERG component

There is considerable trial-to-trial variability in maximum constriction velocity. We sorted trials into 10 equal bins based on maximum constriction velocity, separately for each combination of participant, dataset, and stimulation condition (Fig. 2b). This revealed that the constriction-locked ERG component is most pronounced for trials with a high maximum constriction velocity—exactly as expected, given that a higher pupil-constriction velocity results in a more abrupt decrease in retinal exposure.

#### The constriction-locked ERG component is consistent between datasets

Despite large differences in the shape of the overall ERG response between datasets, all datasets show a positive deflection following constriction (Fig. 2c). The negative deflection preceding constriction is only evident in three datasets.

#### The constriction-locked ERG component originates from the eye electrodes

For the five datasets with complete EEG data, for each electrode separately, a permutation test as described above was performed. The absolute difference between the actual signal and permuted baselines was converted to a z-score effect size (visualized as a scalp distribution in Fig. 2a). This shows that the constriction-locked effect in the 0 - 40 ms constriction-locked window originates from the eye electrodes and radiates out towards nearby frontal electrodes (likely through volume conduction). This pattern strongly suggests a retinal origin of the component.

#### The constriction-locked ERG component is followed by a constriction-locked ERP component over occipital electrodes

For the seven datasets with occipital EEG data, for the average of the occipital electrodes (O1, O2, and Oz), a permutation test as described above was performed. This revealed a positive constriction-locked component around 120 ms after the timepoint of maximum constriction velocity (Fig. 2d). This component is qualitatively different from the constriction-locked component in the ERG (and neighboring frontal electrodes), suggesting that it may reflect a distinct constriction-locked response in visual cortex.

#### Control analyses

The *Supplementary Materials* contain control analyses showing that the constriction-locked ERG component does not result from blinks, eye movements, or muscle artifacts, and a ‘recovery’ analysis to validate the analysis pipeline with simulated data.

### Descriptive ERG results

Fig. 3 shows the ERG signal locked to stimulus onset. A typical ERG response is characterized by a negative component that (for stimulation parameters in our lab; responses are typically faster in clinical ophthalmological settings) peaks around 25 ms after stimulus onset (a-wave) followed by a positive component that peaks around 60 ms after stimulus onset (b-wave). These components are highly sensitive to stimulus intensity (as shown for the Original dataset in Fig. 3b). Most datasets show this typical pattern, with the exception of two datasets in which a dark stimulus was presented against a bright background (Attention-1-bright and Attention-2-bright) which does not elicit an a-wave (but does trigger a slight pupil constriction [Fig. 4] and an associated constriction-locked ERG component [Fig. 2]). Following the b-wave, the ERG signal becomes highly idiosyncratic and variable between datasets. The constriction-locked component is not visible when the signal is locked to stimulus onset, because it is masked by trial-to-trial variability in pupil-constriction latency.

**Figure 4.**
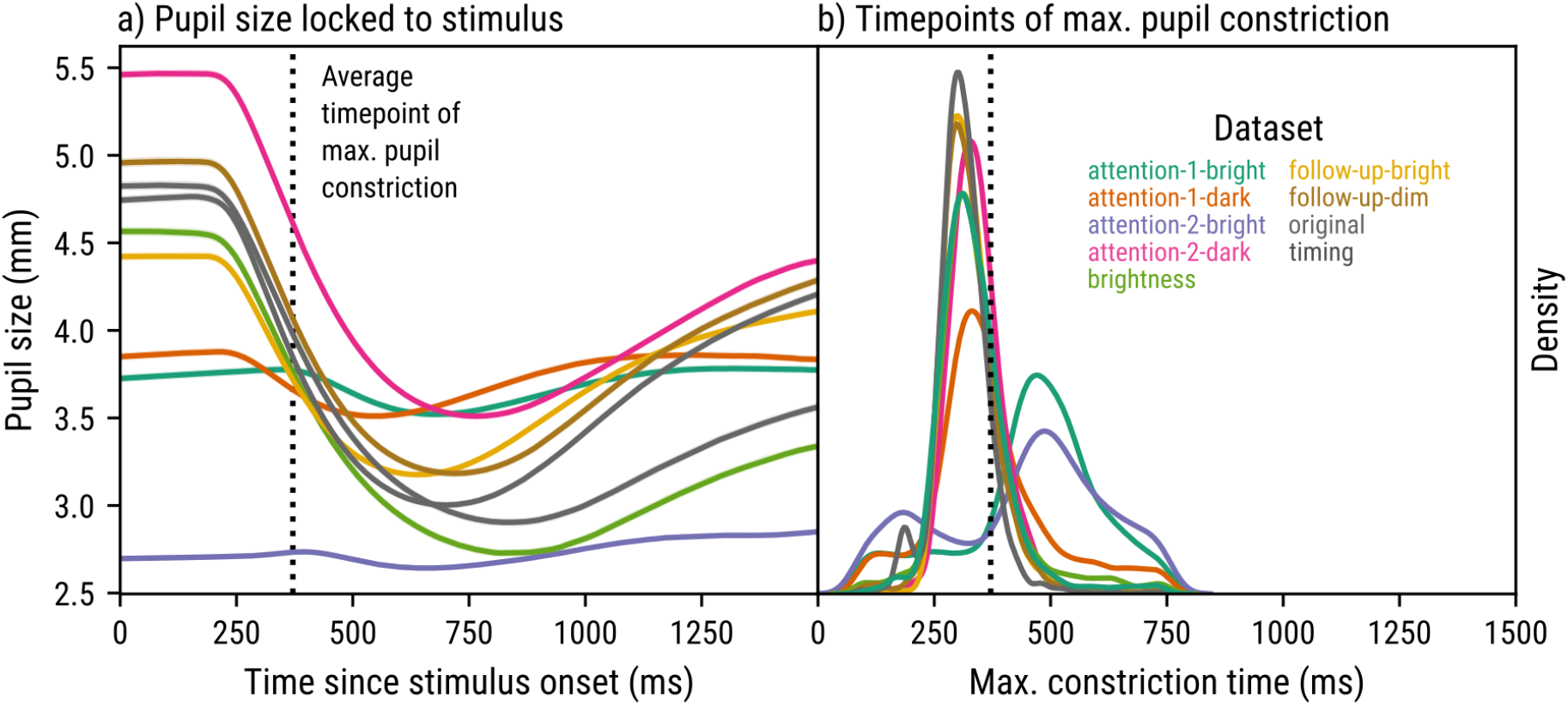
a) Pupil size locked to stimulus onset for all datasets. The vertical dotted line indicates the average timepoint of maximum constriction velocity. b) The distribution (kernel density estimation) of timepoints of maximum constriction velocity for all datasets.

### Descriptive pupil-size results

In all datasets, the pupil started to constrict in response to the stimulus after around 220 - 420 ms, where datasets with strong stimuli (e.g. Original) show a stronger and faster constriction than datasets with weaker stimuli (e.g. Attention-2-bright). The average timepoint of maximum constriction was 371 ms.

## Discussion

Here we show that the human retina and visual cortex respond to pupil constriction, independently of visual stimulation. Most likely, the retina responds to the sudden decrease in retinal light exposure that accompanies pupil constriction, as previously reported for retinal ganglion cells in mice (16). This retinal response then propagates to visual cortex, where it manifests as an event related potential (ERP) that emerges over occipital electrodes around 100 ms after the moment of maximum constriction velocity (roughly 400 ms before the pupil reaches its minimum size).

These findings raise important questions about how the visual system maintains a stable perception of brightness. If the retina responds to changes in light exposure caused by pupil-size changes, why do we not experience corresponding changes in perceived brightness (but see 16 for preliminary psychophysical evidence suggesting that in some conditions pupil constriction does have perceptual effects)? This is largely an open question, but an important clue may come from two recent studies on pupil size and brightness perception (26,27).

Sulutvedt and colleagues (26) asked human participants to judge the brightness of a tester stimulus presented to one eye, relative to a reference stimulus presented to the other eye. The pupil of one eye was pharmacologically dilated, while the other eye was left untreated (and thus had a smaller pupil). Their main finding was that the brightness of stimuli presented to the dilated eye was overestimated relative to those presented to the untreated eye. This finding contrasts with a study by Wardhani and colleagues (27), who asked human participants to judge the brightness of a tester stimulus, relative to a reference stimulus presented earlier (both stimuli were presented to both eyes). In between the presentation of the tester and referent, pupil size was manipulated by presenting red/ blue inducer stimuli (blue stimuli induce sustained pupil constriction; 6,28,29) or by varying cognitive load (high cognitive load induces pupil dilation; 30,31). In this case, increased pupil size did *not* lead to an overestimation of brightness (in fact, there was a slight effect in the opposite direction). An important difference between these two studies is that Sulutvedt and colleagues (26) used a pharmacological manipulation of pupil size, which disrupts the ability of the visual system to monitor pupil size, whereas Wardhani and colleagues (27) used non-pharmacological manipulations, which leave this ability intact. The contrast between these findings therefore suggests that the visual system may be taking pupil size into account when interpreting visual input. Such a mechanism would be consistent with the phenomenon of brightness constancy, where the perceived brightness of objects remains relatively stable despite changes in the amount of light reaching the eye (32). One possible mechanism for achieving brightness constancy may be analogous to models of visual stability with respect to eye movements, which posit that visual input is integrated with a corollary discharge (or efference copy) of the motor commands for eye movements (33,34). However, while the existence of a corollary discharge for eye movements is well-established (35), it is unclear if a similar corollary discharge exists for pupil constriction and dilation. However, it is a theoretically plausible and testable prediction.

Although constriction-induced neural responses pose a challenge to brightness constancy, they may also provide a way to optimize visual perception based on the demands on the current situation (36). For example, pupil dilation (i.e. moments of increasing pupil size, independent of absolute pupil size) is associated with an increased signal-to-noise ratio of activity in mouse visual cortex (i.e. sharper tuning of neural responses; 35). Such findings are often interpreted in terms of arousal-related signals originating from brain areas such as the locus coeruleus. However, it is also possible that pupil dilation itself triggers visual responses in the retina (just as we report here for pupil constriction), which interact with visual processing in a way that enhances the signal-to-noise ratio of visual processing. Most likely, there are multiple routes through which arousal affects visual processing: indirectly through pupil-size changes (38), and directly by modulating activity throughout the visual system (2,39), possibly all the way down to the retina (40–43). An important avenue for future research will be to understand whether and how pupil constriction and pupil dilation induce retinal responses that serve to optimize visual perception.

We have conducted extensive control analyses to verify our claim that the constriction-locked ERG response that we report here indeed originates from the retina and is not an artifact of blinks or eye movements. A further alternative to consider is muscle activity associated with the pupil response itself. Pupil constriction is produced by contraction of the iris sphincter, accompanied by relaxation of the dilator. Although iris tissue is electrically excitable (44,45), there is no evidence that these non-striated muscles generate electromyographic (EMG) activity that can be recorded non-invasively with skin electrodes. The small muscle mass of the iris, its location within the eyeball, and the ring-like geometry of the sphincter make a substantial iris-muscle contribution to the present signal unlikely. Nevertheless, in a dataset of this size, even very weak sources might become detectable, and we therefore cannot definitively rule out contributions of iris muscle activity to our results.

In sum, we report a previously unknown response in the human retina and visual cortex that is locked to pupil constriction, and likely reflects a retinal response to a sudden decrease in retinal light exposure. This finding has important implications for the role of pupil size in visual perception.

